# Differential sensing by the *C. albicans* Gpr1 receptor results in morphogenesis, β-glucan masking and survival in macrophages

**DOI:** 10.1101/2022.11.30.518566

**Authors:** Wouter Van Genechten, Stefanie Wijnants, Jolien Vreys, Patrick Van Dijck

**Author notes:** Patrick Van Dijck, **Email:**. **Author Contributions:** Conceptualization, W.V.G and P.V.D.; Methodology, W.V.G and P.V.D.; Investigation, W.V.G., S.W. and J.V.; Writing – Original Draft, W.V.G.; Writing – Review & Editing, S.W., J.V. and P.V.D.; Funding Acquisition, W.V.G and P.V.D.; Resources, W.V.G., S.W. and J.V.; Supervision, P.V.D.

## Abstract

The human fungal pathogen, *Candida albicans*, is very proficient at several classical virulence factors such as morphogenesis, adhesion, biofilm formation and immune evasion through β-glucan masking. The protein kinase A (PKA) pathway is involved in both morphogenesis and β-glucan masking. Several signals converge onto the PKA pathway, but it contains only a single upstream G-protein coupled receptor, Gpr1. We identified specific residues within the N-terminal tail of Gpr1 that are required for methionine-induced morphogenesis through Tpk2. Furthermore, we observe that Gpr1-Gpa2 has an active role in exposing glucans. Even though Gpr1 is required for survival when *C. albicans* is challenged with macrophages, specifically disrupting morphogenesis did not attenuate this survival. Additionally, constitutive β-glucan masking did not improve *C. albicans* survival rates in the macrophage assay. Taken together, this indicates that Gpr1 may regulate additional mechanisms, possibly through glutamine 461, which are crucial in a macrophage context.

**Significance Statement:** *Candida albicans* is a human fungal pathogen mostly present as a commensal in the gastrointestinal tract. It can rapidly adapt to its everchanging environment through continuous monitoring of extracellular signals. These extracellular signals include methionine and lactate which induce respectively morphogenesis and β-glucan masking through the G-protein coupled receptor, Gpr1. Through a mutagenic approach we different amino acids of the receptor sense methionine and/or lactate but we show that Gpr1 may have an additional ligand that affect its survival in macrophages.

## Introduction

The human fungal pathogen, *Candida albicans*, is highly proficient in adapting to its immediate surroundings which range from the skin to mucosal surfaces and blood (1–3). It can be appreciated that these host infection niches all differ widely in their static or fluctuating nutrient concentrations. Therefore, nutrient sensing is of utmost importance to *C. albicans*. To constantly adapt to the rapidly changing environment it has a wide set of sensors for physical and nutritional stimuli. Generally, sensing is mostly performed by G-protein Coupled Receptors (GPCRs) which are found across the entire eukaryotic kingdom and are classically divided in six classes (4). An additional seventh class is proposed, that of the fungal nutrient receptors, which is further divided in ten subclasses. The third subclass, being the Rhodopsin-like/carbon subclass, of these fungal nutrient receptors, contains receptors with a high homology to the glucose and sucrose sensing *S. cerevisiae* Gpr1. This class contains the *C. albicans* Gpr1 which was first discovered in a screening of genes that are upregulated in response to phagocytosis by macrophages (5). A year later, Miwa and colleagues discovered the role of Gpr1 in the Protein Kinase A (PKA) pathway and subsequent filamentation on solid media (6). Nevertheless, contradicting results were obtained concerning the role of Gpr1 in cAMP accumulation. Maidan and colleagues describe that accumulation of cAMP after stimulation with glucose or serum is dependent on the Cdc25-Ras1 module and not on the Gpr1-Gpa2 module as stated by Miwa and colleagues (7, 8). Additionally, whilst the virulence of a homozygous *GPR1* deletion strain is attenuated in a systemic mouse infection model by Maidan *et al*., no such virulence defect was observed for a *GPR1* or *GPA2* deletion in a similar lateral tail vein infection model by Miwa and colleagues (7, 8).

Since glucose is not involved in the Gpr1-Gpa2 mediated cAMP increase in *C. albicans*, another component of the hyphae-inducing media must be the ligand of Gpr1. It was discovered that amino acids are involved since addition of alanine or methionine leads to a rapid internalization of Gpr1, a common reaction to the ligand-receptor interaction, which activates the yeast-to-hyphae transition on solid media and tissue models. Additionally, hyphae formation on solid media was successfully achieved by supplementing synthetic minimal medium with proline or methionine in the presence of low concentrations of fermentable carbon sources such as glucose, fructose, maltose, sucrose or galactose (7). However, relatively high non-physiological concentrations of glucose repress filamentation, indicating that there is crosstalk between the multiple signals and that the nutrients must be sensed and/or transported at certain concentrations to fully activate PKA signaling and filamentation (9). Indeed, methionine-induced hyphae formation through Gpr1 and the cAMP-PKA pathway also requires the transport of methionine by the high affinity transporter Mup1 and subsequent metabolization (10). Comparable to the *GPR1* deletion, a *MUP1* deletion (or the conditionally-repressed *SPE2*, encoding S-adenosyl methionine decarboxylase (SAM) that is converting SAM into S-adenosylmethioninamine, a precursor in polyamine biosynthesis) results in an attenuated and delayed virulence in a murine systemic infection model (11).

Recently an additional role of Gpr1 signaling was discovered. Lactate can trigger β-glucan masking within the cell wall. β-glucan masking is an immune evasion strategy where the pathogen-associated molecular pattern (PAMP), cell wall carbohydrates such as β-glucan, are masked within inner layers of the cell wall, with more exposure of less immunogenic (for macrophages) mannoproteins. This masking limits recognition of the PAMP by Dectin-1, the pattern recognition receptor (PRR) of the host immune system. It was shown that this masking depends on Gpr1, but β-glucan masking was not significantly affected in mutants of the PKA pathway. Therefore, this signaling event was proposed to be completely independent of the cAMP-PKA pathway and its role in filamentation. The novel signaling function of Gpr1 was proposed to operate via a putative second Gα subunit, Cag1, which they suggested to inhibit the masking (12). Additionally, the transcription factor Crz1, which functions downstream in the calcineurin pathway, was shown to be involved in the masking process. This illustrates that this novel pathway utilizes components, Gpr1, Cag1 and Crz1, from different well-characterized pathways (12). Nevertheless, recent papers report a role of the PKA pathway in β-glucan masking through Sin3 and Mig1/Mig2 regulating exo- and endoglucanases (13).

Currently, it is broadly established that GPCRs can have multiple ligands and interact with several G proteins or even that several GPCRs can share a G protein (14). Therefore, it is not unreasonable that Gpr1 can have methionine and lactate as ligands. Furthermore, Ballou and colleagues hinted at the similarity in structure between methionine and lactate and that they may have overlapping ligand binding sites on Gpr1 (12). Furthermore, other short chain fatty acids such as acetate and butyrate may serve as antagonists of Gpr1-Gpa2-induced β-glucan masking (15).

We further characterized the different roles of Gpr1 through a mutagenic approach. Specific point mutants were created that allowed us to investigate the different roles of Gpr1 and its downstream effects. We confirmed (and therefore contradict the Ballou et al. hypothesis) that there is a single Gα protein, Gpa2, associated with Gpr1 and that the Gpr1-Gpa2 module activates morphogenesis as well as lactate-induced glucan masking. Whereas the former phenotype is mediated solely by the cAMP-PKA pathway (10), the latter phenotype has both PKA dependent and independent components. Indeed, β-glucan masking is partly regulated by the Tpk isomers but this is independent of sensing lactate and most likely acts through regulation of the mannan layer covering the β-glucan layer as was proposed elsewhere (16, 17).

Furthermore, we evaluated these Gpr1 point mutants and their respective phenotypic defects upon interaction with macrophages. We conclude that neither immune evasion nor amino acid induced filamentation determine the outcome and fitness of *C. albicans* in a macrophage survival assay, indicating that another downstream phenotypic effect of Gpr1 plays a major role in macrophage survival. We hypothesize that the main determinant of survival of *C. albicans* when confronted with macrophages is stress resistance which is regulated through a putative G_β_ protein.

## Results

### Alignment of *Ca*Gpr1 results in the selection of several important residues for site-directed mutagenesis

Since fungal GPCRs form an independent class, they are highly homologous and contain conserved residues for sensing and signaling that may bind common ligand substructures. Alignment of the Gpr1 amino acid sequences of *C. albicans* and *S. cerevisiae* (Fig. 1A), of which residues important for sucrose and glucose signaling have already been identified (18), is an ideal starting point for a mutagenic analysis. The residues selected are tryptophan 451 and glutamine 461 as indicated in figure 1A. Tryptophan 451 is situated in the middle of the sixth transmembrane domain, whilst glutamine 461 is located just extracellularly and should therefore be accessible to possible ligands. The N-terminal extracellular tail of *Ca*Gpr1 is significantly longer compared to that of *Sc*Gpr1, possibly indicating that this has an additional role in nutrient sensing in *C. albicans*. In the eukaryotic world, such extensive N-terminal tails in GPCRs have been reported in mammalian taste receptors. We therefore also aligned the N-terminal tail of *Ca*Gpr1 with several glutamate and taste receptors as shown in figure 1A. Even though sequence identity between these sequences is lower, we were able to identify two conserved residues, serine 34 and threonine 56. A three-dimensional model of Gpr1 that was predicted with Alphafold is depicted in figure 1 panel B and C (19, 20). These panels show the location of the selected residues on a side view of the entire protein (Fig. 1C) where the long extracellular highly unstructured tail is evident and serine 34 and threonine 56 are present in two putative a-helices (orange), which are predicted with a low confidence (50 < per-residue confidence score (pLDDT) < 70). In panel 1D a closer view of the transmembrane helices, which have a very high confidence score (pLDDT>90), is depicted (blue).

**Figure 1.**
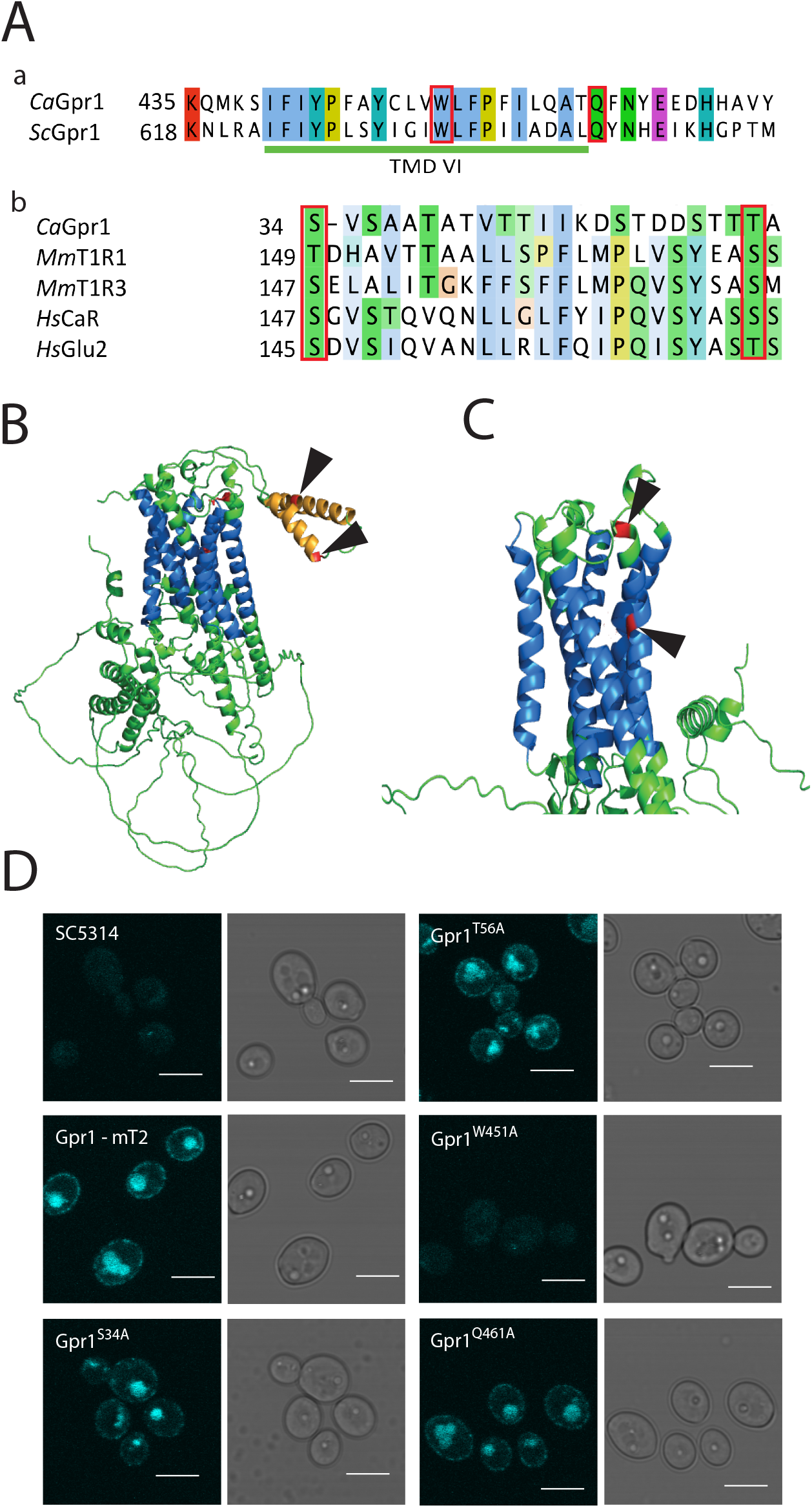
Alignment of CaGpr1 results in the selection of several important residues for site-directed mutagenesis. A) Alignment of CaGpr1 with a) the TMDVI of ScGpr1 and b) the N-terminal tail of mammalian taste receptors. Colours indicate the charge of the residue with red denoting a positive charge, magenta a negative charge and polar residues in green. B) Cartoon scheme of the three-dimensional structure of Gpr1 as predicted by alphafold. Arrows indicate the S34 and T56 residues in the N-terminal tail. C) Closer look at the 7 transmembrane domains. Arrows indicate the W451 and Q461 residues. D) Confocal fluorescence microscopy of mutated Gpr1 fused with mTurquoise2. Scalebar is 5 μM.

The residues selected based on the alignment as indicated in figure 1 were mutated to alanine using site-directed mutagenesis. Furthermore, we overexpressed the mutated Gpr1 in a fusion construct with mTurquoise2 to assess expression levels and correct localization. In figure 1D we can see a correct membrane localization pattern of the wild type Gpr1–mTurquoise2 fusion. Some intracellular mTurquoise2 signal is observed, possibly as a result of the overexpression which may induce rapid turnover and internalization of the GPCR (21). Furthermore, mTurquoise2 fusion constructs with Gpr1 mutated at S34, T56 or Q461 mimic the membrane localization pattern. This indicates that these strains can be used for further phenotypic analysis. In contrast, mutating W451, which is situated within transmembrane domain VI, abolished the signal completely. We can therefore conclude that there is no noticeable expression of this construct. We also constructed combinations of the S34, T56 and Q461 mutations which are depicted in supplementary figure 1.

### The N-terminal tail of Gpr1 is required for the methionine-induced morphogenesis and requires Gpa2 for signaling

Morphogenesis is one of the major virulence factors of *C. albicans* and one possible way to induce filamentation is through methionine-induced activation of the PKA pathway via the Gpr1-Gpa2 sensing system. Activation of this pathway may occur during tissue infection upon digestion of proteins through Secreted Aspartyl Proteases (SAPs) or within the bloodstream. Using the point mutants that we generated in Gpr1 we aimed to unravel which amino acids/domains in Gpr1 are involved in methionine sensing *in vitro*. First, we confirmed that the wild type SC5314 displayed extensive methionine-induced filamentation, whilst filamentation is almost completely abolished in the *GPR1* deletion strain (Fig. 2A). Reintegrating wild type *GPR1* or fused with mTurquoise2 restored the filamentation defect of the *GPR1* deletion strain. Furthermore, it is clear from figure 2A that residues within the N-terminal tail, S34 and T56, are required for methionine-induced filamentation. On the other hand, Q461 is not involved in methionine signaling. As expected from the localization assay, W451 showed the same phenotype as the *GPR1* deletion strain since no expression of the construct was visible. Activation of filamentation by methionine sensing occurred solely through Gpa2 and not by the putative Ga protein, Cag1 (Fig. 2B). Morphogenesis data for the double and triple point mutants are found in supplementary figure 2. As expected, double and triple mutants have attenuated or abolished morphogenesis since either mutating the S34 or T56 is sufficient to disrupt methionine signaling.

**Figure 2.**
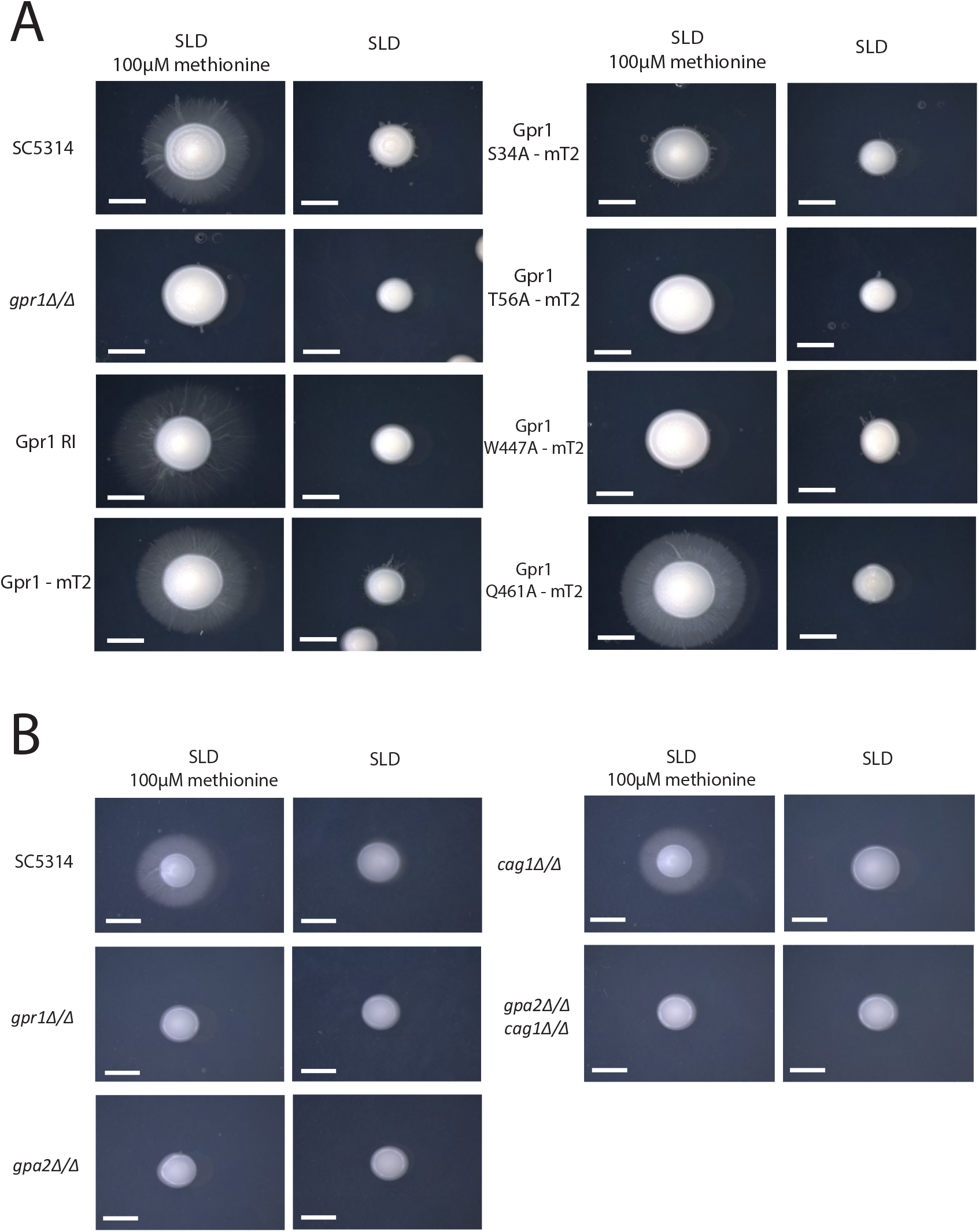
The N-terminal tail is required for the methionine-induced morphogenesis through Gpa2. Morphogenesis assay on solid medium with and without 100μM L-methionine. Direct comparison of the SC5314 and GPR1 deletion strain with A) the Gpr1 point mutants and B) the putative Gα subunits. A representative colony is depicted for each strain in each condition. The experiment was performed twice with three biological repeats. Scalebar denotes 5 mm.

### Gpr1, through its N-terminal tail, and Gpa2 have an active role in exposing glucans

Previously, it was shown by Ballou et al. that Gpr1 is required for lactate-induced masking of the glucan layer in the cell wall, which is an immune evasion strategy (12). Here we can confirm that lactate sensing results in a reduced exposure of β-1,3 glucans in the wild type SC5314 strain (Fig. 3A). However, in contrast to what was reported by Ballou and colleagues, the *GPR1* deletion strain showed constitutively low β-1,3 glucan exposure in both glucose and glucose + lactate grown conditions. The N-terminal S34A mutant showed attenuated masking whilst the T56A and Q461A mutants showed a wild type phenotype with significant masking (p=0.0329 and p=0,0042 respectively) (Fig. 3A). The W451A mutant highly resembled the *GPR1* deletion strain with constitutive masking in both glucose and glucose + lactate grown conditions. Interestingly, from figure 3B we can see that the glucan masking effect is also attenuated through Gpa2 and not the putative Ga protein, Cag1, which still showed significant masking upon administration of lactate (p<0,0001). The β-1,3 glucan masking of the double and triple point mutants is depicted in supplementary figure 3. All double and triple mutants showed defects in lactate-induced β-1,3 glucan masking.

**Figure 3.**
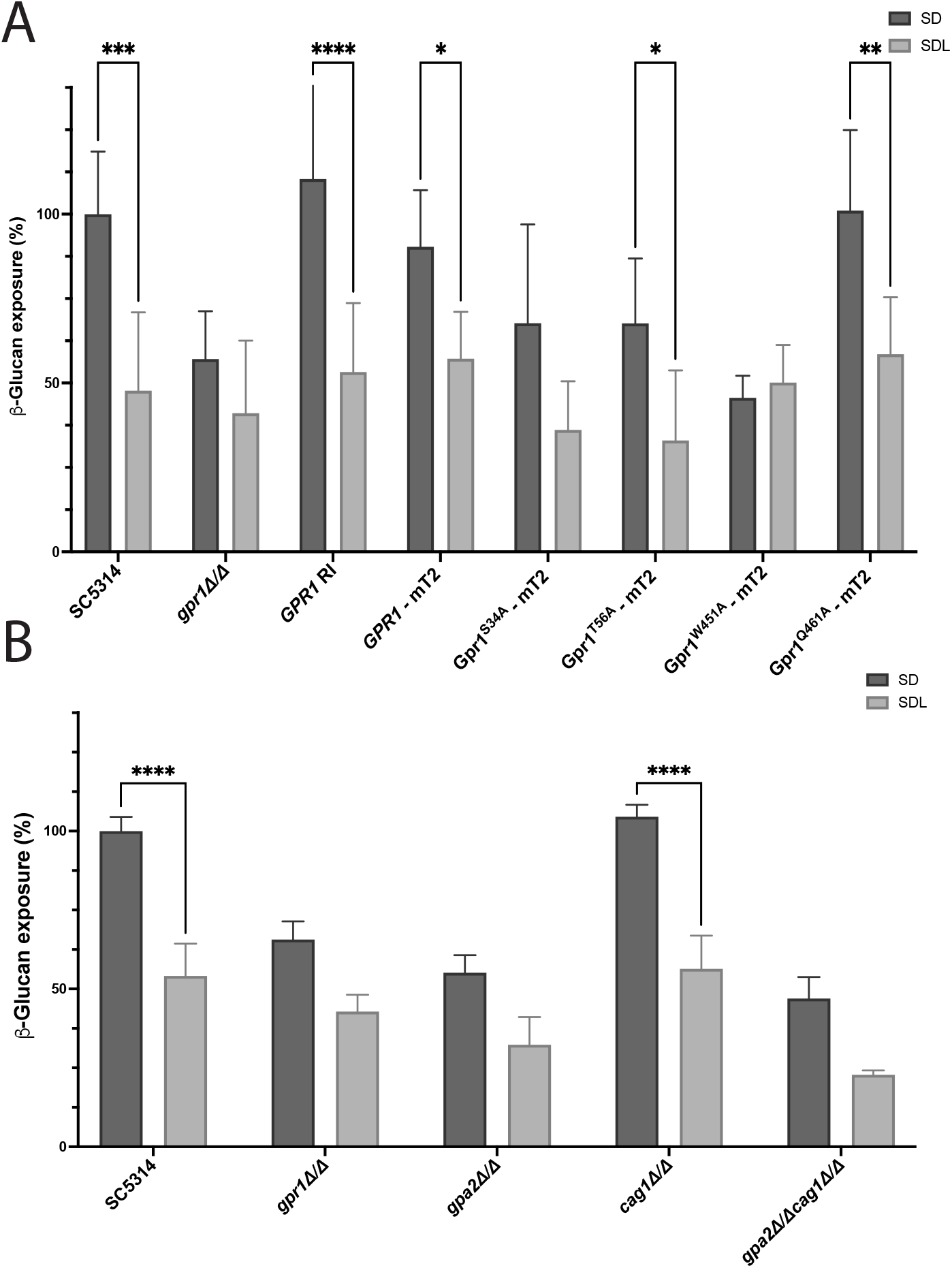
Gpr1, through its N-terminal tail and Gpa2, has an active role in exposing glucans. Exposure of □-glucan as measured with flow cytometry after labeling with alexa488 utilizing hDectin1. Median fluorescence intensity of 5 000 cells is normalized to SC5314 in SD conditions. A) Analysis of masking of the Gpr1 point mutants. B) Assessment of masking of the GPA2 and CAG1 deletion strains upon administration of 2% lactate. Statistical analysis was performed using two-way ANOVA with Sidak multiple comparison correction. 3 biological repeats were utilized for each strain in each condition and the experiment was performed twice. Asterisks denote significance levels with; *, P < 0.05; **, P < 0.01; ***, P < 0.001; ****, P < 0.0001. Depicted error bars show the mean and SEM.

### Gpr1 is required for increased survival when *C. albicans* is challenged with macrophages

Immune evasion through epitope masking result in decreased recognition by host immune cells, such as macrophages which should result in increased survival of *C. albicans* when challenged with macrophages (13, 22). However, in our hands, deletion of *GPR1* results in a significantly lower survival rate (p = 0,0001) compared to SC5314 in such a macrophage co-culture assay (Fig. 4A). Reintegrating *GPR1* restored the survival percentage to the wild type level. The Gpr1 strains containing the S34A and T56A mutations did not result in a significant difference in survival rate compared to SC5314. In contrast, the W451 (p=0,0338) and Q461 (p=0,0108) mutation did show significantly attenuated survival. Supplementary figure 4 depicts the double and triple point mutants. The double point mutant containing both the S34A and T56A did not influence survival in the presence of macrophages, whilst mutating glutamine 461 did result in significantly lower survival of *C. albicans* in this assay. As shown in figure 4B, deleting *CAG1* did not influence survival of *C. albicans* in the presence of macrophages, however deletion of *GPA2* increases the survival rate significantly (p=0,0173). The double deletion strain, *gpa2Δ/Δ cag1Δ/Δ* has a wild type survival rate. Since we establish that both methionine-induced filamentation and lactate-induced β-glucan masking is regulated through Gpa2, we investigated whether the signal would be diverged further downstream the cAMP-PKA pathway.

**Figure 4.**
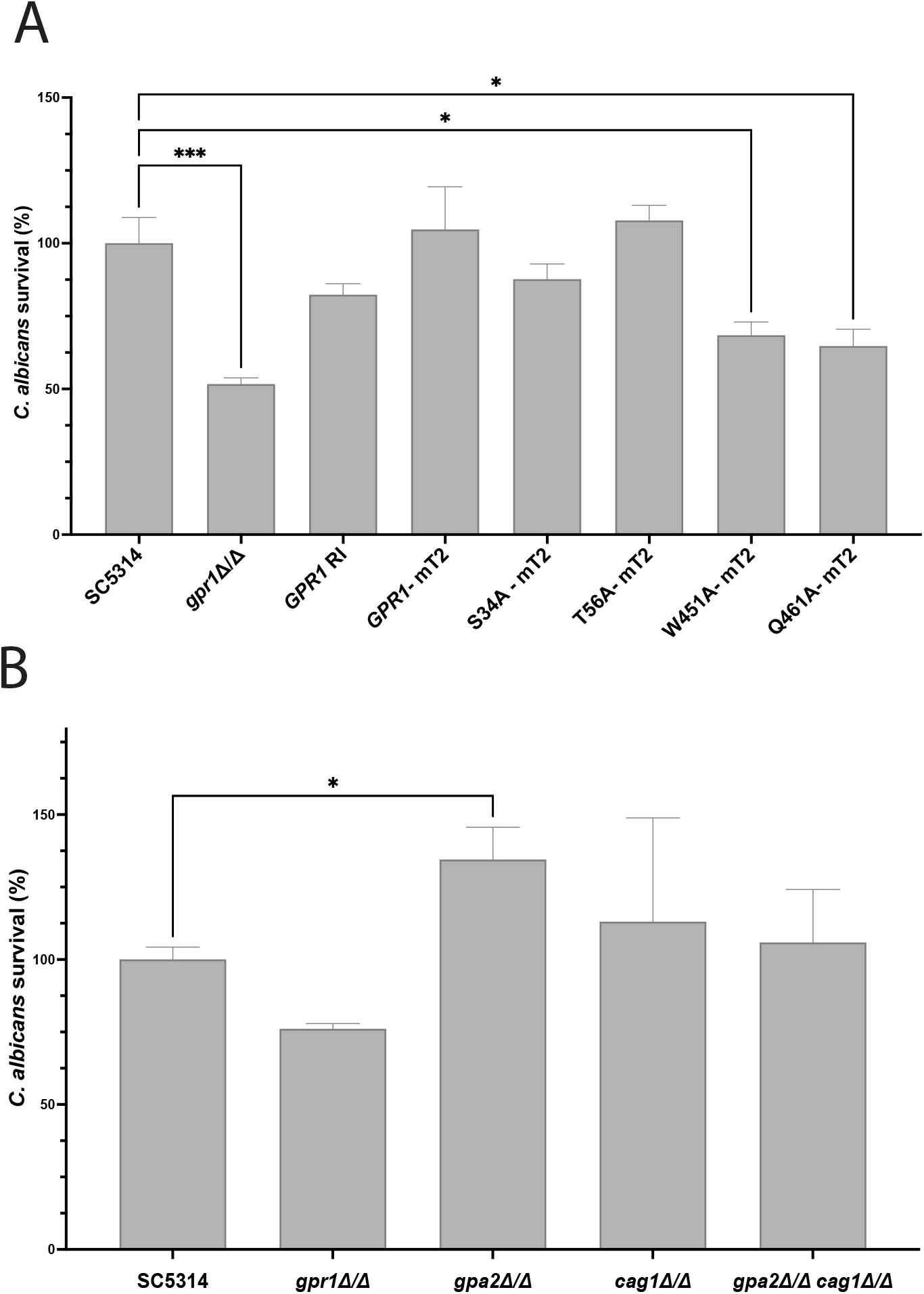
Gpr1 is required for increased survival when C. albicans is challenged with macrophages. Candida albicans survival assay when co-cultured with BMDMs for 3h at 37°C. Fungal cells were plated and counted to score for survival. A) Survival rate of the Gpr1 point mutants. B) Survival rate of the GPA2 and CAG1 deletion strains. Results shown are the average of three independent experiments containing three experimental repeats of a single biological repeat. Raw data is normalized to the survival rate of the wild type SC5314. Statistical analysis was performed using one-way ANOVA with a Bonferroni-corrected multiple comparison to the wild type. Asterisks denote significance levels with; *, P < 0.05; **, P < 0.01; ***, P < 0.001; ****, P < 0.0001. Depicted error bars show the mean and SEM.

### Both PKA catalytic subunits differentially regulate glucan exposure while only Tpk2 is required for methionine-induced morphogenesis

Earlier reports described differential roles of the two PKA catalytic isoforms, Tpk1 and Tpk2. In general, Tpk2 shows a higher kinase activity compared to Tpk1 and both isoforms are required for morphogenesis in liquid and solid media under certain conditions (23). On solid medium supplemented with methionine, filamentation and agar invasion were completely abolished in the *tpk2Δ/Δ* strain whilst it was not attenuated in the *tpk1Δ/Δ* strain, indicating that Tpk2 is required for methionine-induced filamentation and that the remaining isoform Tpk1 has insufficient activity to support the induction of filamentation under methionine-supplemented conditions (Fig. 5A). In contrast, deletion of *TPK1* displays a constitutively lower glucan exposure of the cells in the glucose grown condition, even before addition of lactate to stimulate masking (p = 0,0149) (Fig. 5B). Even though the glucan exposure in the glucose condition of the *TPK1* deletion strain is altered, both the *tpk1Δ/Δ* and *tpk2Δ/Δ* significantly masked glucans upon addition of lactate (p = 0,0064, P = 0,0036). A significant difference is observed between the lactate-induced glucan-masked state of the *tpk2Δ/Δ* strain compared to SC5314, where the *tpk2Δ/Δ* strain still displays more glucans (P = 0,0206). This could indicate that part of the masking, through a generic activity of the PKA pathway, is occurring through Tpk2.

**Figure 5.**
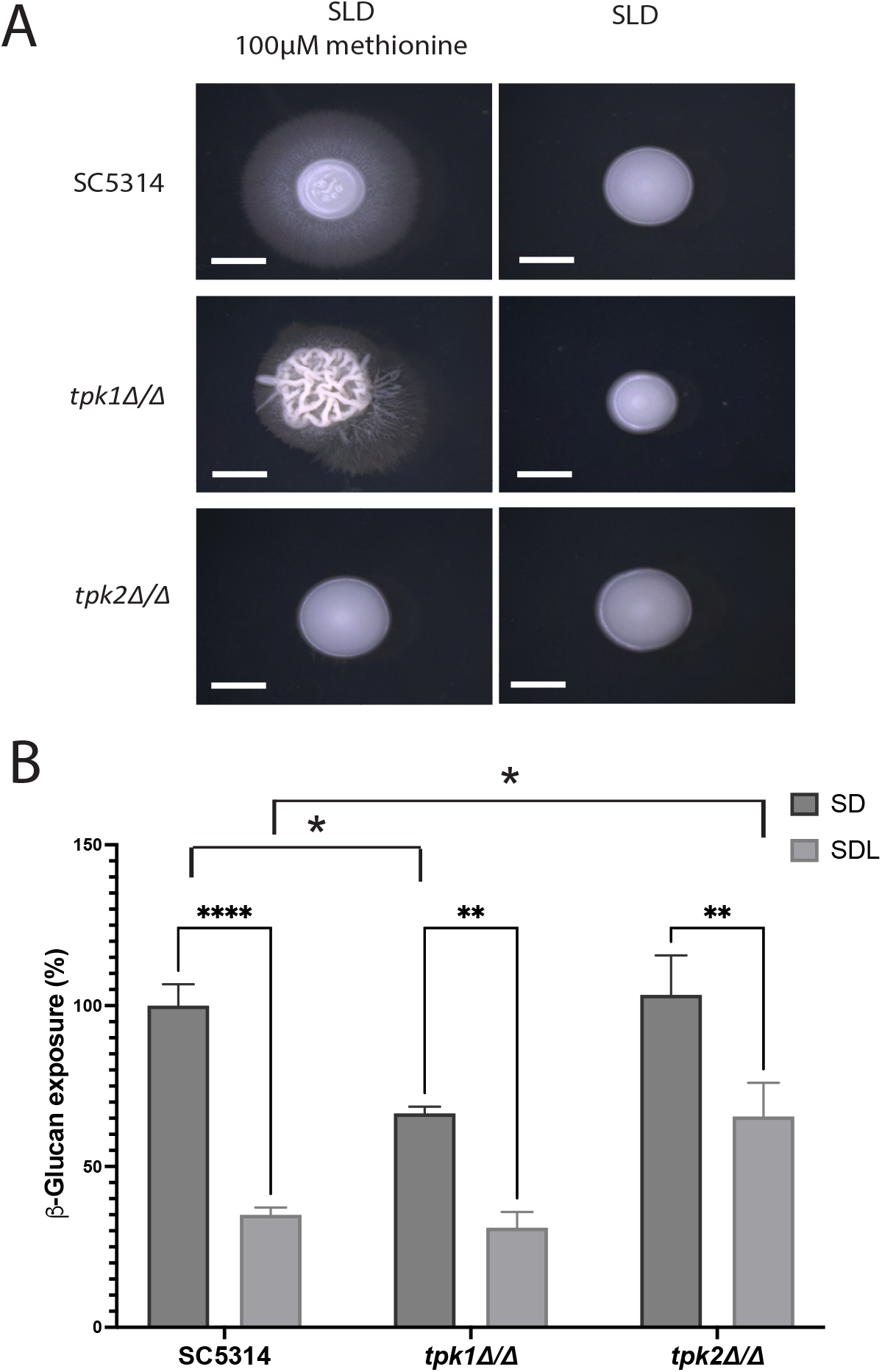
Both PKA catalytic subunits differentially regulate glucan exposure and only Tpk2 is required for methionine-induced morphogenesis. Assessment of A) methionine-induced morphogenesis and B) lactate-induced □-glucan masking of the TPK1 and TPK2 deletion strains to elucidate the role of these isomers within the PKA pathway and downstream phenotypes. The scalebar in the morphogenesis assay (A) denotes 5 mm. Statistical analysis was performed using two-way ANOVA with Sidak multiple comparison correction. 3 biological repeats were utilized for each strain in each condition and the experiment was performed twice. Asterisks denote significance levels with; *, P < 0.05; **, P < 0.01; ***, P < 0.001; ****, P < 0.0001. Depicted error bars show the mean and SEM.

## Discussion

In this work we establish through a mutagenic approach that the methionine binding residues of Gpr1 are present in the extracellular N-terminal tail, specifically S34 and T56. Through a Gpa2-dependent mechanism adenylate cyclase is activated which produces cAMP (24, 25). This cAMP releases the catalytic subunits of PKA, and only Tpk2 is required for the methionine-dependent morphogenesis on solid media. The specific requirement of Tpk2 in this methionine-induced morphogenesis is of great importance since Tpk2 is required for full virulence in both murine systemic and oropharyngeal candidiasis (26, 27). Even though the PKA pathway was shown to have significant effect on *in vivo* virulence, it is not required for filamentation *in vivo* where the transcriptional response inducing hyphae formation is mainly acquired through cAMP-independent pathways (28). Nevertheless, *in vitro* both Tpk isomers were reported to have overlapping and partly redundant roles in growth and morphogenesis with some preferences for inducing ligands and conditions. Whilst Tpk2 is responsible for the majority of phosphotransferase activity, a *TPK2* null mutant is still capable of morphogenesis by induction of N-acetyl glucosamine in liquid (23) or on solid spider or serum supplemented medium (6), indicating that the remaining Tpk1 subunit has sufficient PKA activity. Nevertheless, a distinction was made that Tpk1 displays a larger role in morphogenesis on solid medium compared to Tpk2 and vice versa in liquid conditions (29). Our results partly contradict this observation and further specify that one of the ligands responsible for hyphal induction within serum is methionine. Methionine and other free amino acids are relatively abundant in plasma with methionine being reported reaching concentrations of approximately 30 and 64 μM in respectively humans and rats (30, 31). Furthermore, in the host, other signals such as an ambient temperature of 37°C also stimulates morphogenesis. The methionine-induced morphogenesis is still present when plates were incubated at 37°C, although there is also some filamentation in the conditions without any supplemented methionine. This is due to the well-known temperature-dependent filamentation which is Gpr1 independent (32).

The second phenotype that is controlled through Gpr1 is β-glucan masking, which is also aptly called epitope shaving since Xog1, a secreted exoglucanase, removes β-glucan from the exposed cell surface (33). We report that a *GPR1* deletion strain shows constitutively lower β-glucan exposure, indicating a negative regulation by Gpr1. Reintegrating *GPR1* restores the initial β-glucan exposure of the cells when grown in regular glucose-containing conditions. The N-terminal residues, S34 and T56, which were responsible for methionine-induced filamentation also show a decreased β-glucan exposure in glucose-grown conditions. However, mutation of S34 or T56 has only a slight effect on β-glucan masking upon lactate addition compared to the *GPR1* re-integrants or the Gpr1^Q461A^ mutant. This indicates that activation of masking through Gpr1 occurs independently of these N-terminal residues. In contrast to the report of Ballou and colleagues, not the putative G protein, Cag1, but Gpa2 is required for masking (12). These observations suggest that the conventional PKA pathway is involved in cell wall remodeling, but only Gpr1 and Gpa2 are required for the lactate-induced masking. Similar to the investigation of the Tpk isomers in morphogenesis, we investigated the role of these two catalytic subunits in β-glucan masking (Figure 5). To our surprise, the two catalytic subunits do not affect the lactate-induced β-glucan masking but have contradicting roles in epitope exposure in non-lactate, glucose-grown conditions. In the non-lactate condition Tpk1 is required for β-glucan exposure, whilst Tpk2 limits the extent of β-glucan exposure upon addition of lactate. These two catalytic subunits therefore balance the glucan exposure. The resulting phenotype can be acquired in several ways such as the already described epitope shaving through exoglucanases, however exoglucanase activity is low in medium containing glucose (33). Another possibility to limit glucan exposure could be a lower production of glucans in general. However, glucans form the most important structural component and serve as an anchoring point of glycoproteins. Therefore, we assume that this process of immune evasion that would be detrimental to the strength of the cell wall is not advantageous. Furthermore, there is no correlation between the absolute quantity of β-glucans and the exposure of these glucans (22, 33, 34). Finally, another possibility to mask the glucans is by increasing the top layer of mannans and proteins (35). Although mannans themselves are a PAMP, the immune response to mannans is dependent on cell type and specifically macrophages have a lower cytokine response to mannans compared to β-glucans (36). Therefore, in addition to the lactate-induced β-glucan masking independent of the PKA pathway, we hypothesize that the PKA catalytic subunits regulate, in opposite manners, the mannan production to balance excess mannan production and macrophage recognition through β-glucans.

The environmental lactate which triggers this PAMP masking may originate from the microbiota (e.g., it can be produced by lactic acid bacteria) or from the host (e.g., by macrophages), possibly linking fungi, bacteria, and the immune system in a dynamic complex. Furthermore, Gpr1 is upregulated during phagocytosis (5) indicating that *C. albicans* has ample availability to sense ligands which may result in immune evasion or limitation of uptake through hyphal morphogenesis. We confirm the importance of Gpr1 in the interaction with the innate immune system since a *GPR1* deletion strain is severely affected in survival when challenged with macrophages (Fig 4A). Surprisingly, mutating S34 or T56 does not have an effect on *C. albicans* survival, whilst these residues were important for methionine-induced morphogenesis. This indicates that either methionine is not present in sufficient quantity to induce morphogenesis, or that this methionine-induced morphogenesis does not have a beneficial role in survival when challenged with macrophages. If this regulation of morphogenesis is not required, is Gpr1-dependent immune evasion through β-glucan masking important? Although the S34A and T56A mutants have a very slight defect in the lactate-induced β-glucan masking, they do not display an attenuated macrophage survival. Furthermore, the Q461A mutant has a severe macrophage survival defect, but no lactate-induced β-glucan masking defect. These two observations confirm that β-glucan masking is not required for survival of *C. albicans* within this macrophage survival assay. The *TPK1/2* deletion strains were not included within the macrophage assay since they affect several downstream effectors and would not result in a clear dissection of the role of Gpr1 within the fungus-host interaction. Strikingly, we observed that Gpa2 is a negative regulator of survival in the presence of macrophages. This effect is even more surprisingly since this Gpa2 mutant showed a slightly delayed lag phase when grown in the differentiation medium at 37°C which is used during the macrophage assay (Suppl. Fig 6). Furthermore, deleting both *CAG1 GPA2* restores the survival to wild type levels, suggesting that Cag1 acts downstream of Gpa2 in the key factors regulating survival in the macrophage assay.

Since the Q461A mutant has a survival defect and this defect is independent of the morphogenesis and masking assays, a third role of Gpr1 must be present which is regulated through this specific residue. Within macrophages *C. albicans* encounters reactive oxygen and nitrogen species, degradative enzymes (37), iron-deprivation (38),… Therefore, we predict that the additional role of Gpr1 will be the regulation of certain stress responses. Although a general stress response is absent in *C. albicans* (39), the possibility remains that Gpr1 regulates certain stress responses important for survival of macrophage derived stresses. It was already reported before that Gpr1-dependent activation of PKA reduces trehalose biosynthesis and that disrupting *GPR1* leads to trehalose accumulation and suppresses filamentation (40). However, this trehalose accumulation does not fit with the observation that a *GPR1* deletion strain still has a lower survival rate in the macrophage survival assay. The experiments cannot really be compared as both types of experiments were performed in different media with different glucose concentrations. Furthermore, the contrasting roles of Gpr1 and Gpa2 in this assay indicate that Gpr1 may negatively regulate Gpa2 for some phenotypes, possibly through sequestration, membrane localization and/or GDP-GTP exchange inhibition. On the other hand, stimulation of Gpr1 by lactate or methionine can release Gpa2 and stimulate GTPase activity. According to homology with *S. cerevisiae*, we propose the signaling scheme in figure 6, where a putative GTPase binding protein, orf19.4557, has a role similar to the Kelch Repeat Homologues (Krh1/Gpb2, Krh2/Gpb1) in *S. cerevisiae* (41, 42). The gene product of orf19.4557, henceforth called Krh1, is a putative G_β_ protein that interacts with Gpa2. This interaction would be mediated through Gpr1 and rely on the activation state of Gpa2, activation of Gpr1 and subsequent formation of GTP-bound Gpa2 would activate Cyr1 and sequester Krh1. Deleting *GPA2* could result in the release of Krh1, which like *Sc*Krh1 directly inhibits PKA and thus results in stress resistance and increased survival when confronted with macrophages. As mentioned earlier, the Q461 mutant negatively affects the survival rate of *C. albicans* in a macrophage co-culture, independently of glucan masking or morphogenesis. Therefore, this specific residue can be involved in regulating the interaction between Gpa2 and Krh1.

**Figure 6.**
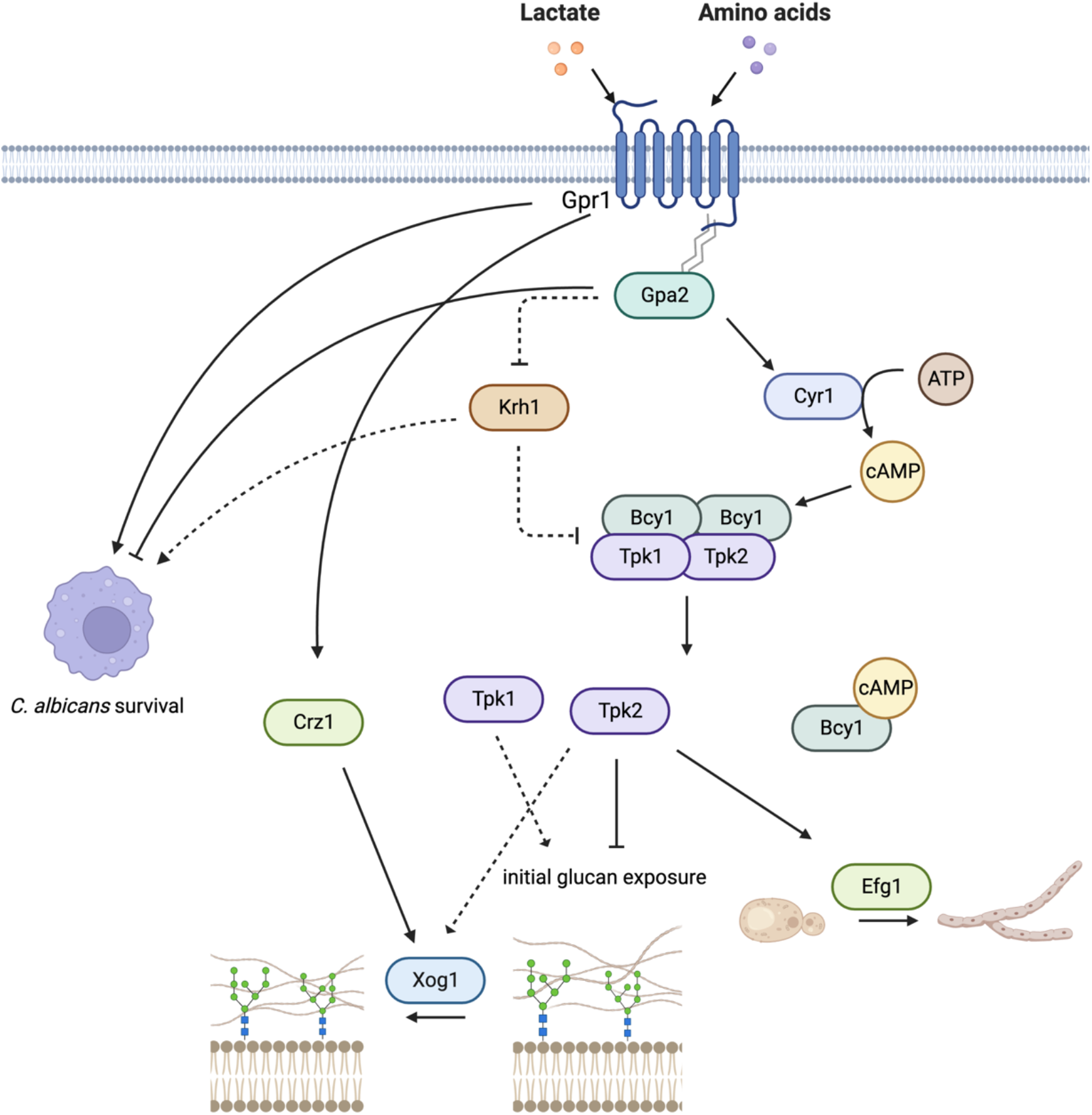
Schematic model of Gpr1 signaling and its implications in morphogenesis, glucan masking and macrophage survival. Hypothetical model linking the Gpr1-Gpa2 module to the PKA pathway through a cAMP-dependent process. This classical PKA pathway is the activator of morphogenesis when supplemented with methionine. The Gpr1-Gpa2 module masks glucans upon sensing lactate through a process independent of cAMP. Furthermore, Gpr1-Gpa2 also regulates glucan masking through Tpk1 and Tpk2, possibly through mannan regulation.

To conclude, Gpr1 is an important sensor for *C. albicans* adaptation to the immediate environment and regulate interkingdom interactions. Methionine promotes filamentation on solid medium through binding with the N-terminal residues S34 and T56 of Gpr1. The Gpr1-Gpa2 module (7) eventually activates specifically Tpk2, not Tpk1. The previously reported lactate-induced β-glucan masking (12) occurs through Gpa2, not Cag1, and is independent of the Tpk isomers. Through a yet undefined pathway Gpr1-Gpa2 will activate Crz1 and subsequently shaves epitopes through the action of the exoglucanase Xog1 (33).

Independent of the lactate-induced immune evasion, Tpk1 and Tpk2 have opposite roles in β-glucan masking, most likely through respectively diminished or enhanced mannan layering. The upstream signals influencing the Tpk activity in this process are unknown, this could be the result of an assembly of upstream signals converging onto the PKA pathway. Neither the β-glucan masking or filamentation on solid media are the main factor of *C. albicans* survival in contact with macrophages. We hypothesize that the main determinant of survival in this assay could be general stress resistance. We propose that a putative homolog of the *S. cerevisiae* Krh1/Gpb2 and Krh2/Gpb1, orf19.4577, may positively regulate stress resistance and thus increase *C. albicans* survival in a macrophage context. The Gpr1-Gpa2 module may inhibit this *C. albicans* Krh1 through sequestration, releasing the inhibition through deletion of *GPA2* leads to higher stress resistance ultimately determining the survival of *C. albicans* in a macrophage context.

## Materials and Methods

### Gene deletion using CRISPR-Cas9

All strains created and utilized in this study are presented in Supplementary table 1. *GPR1, GPA2, CAG1, TPK1* and *TPK2* deletions were created using the CRISPR genome editing method to delete both alleles (43). According to the reported method by Nguyen and colleagues we created gRNA fragments by fusion PCR of a universal A fragment from template pADH110 with a gene specific B fragment. Primers and plasmids utilized to create these fragments are listed in respectively supplementary table 2 and supplementary table 3. The *CAS9* gene is encoded on the pADH99 plasmid and linearized through restriction by MssI. gRNA fragments and linear pADH99 were transformed simultaneously into *C. albicans* and selection was performed with nourseothricin. At least three correct colonies were grown on YP maltose (2%) to induce a flippase and remove the nourseothricin marker and *CAS9*.

### Construction of Gpr1 overexpression strains, mTurquoise2 fusions and specific point-mutations

We reinserted *GPR1* within the *gpr1Δ/Δ* strain using an overexpression construct in a CIp10 where the *URA3* marker was previously replaced by a *NAT1* gene conferring resistance to nourseothricin. This plasmid was digested with high fidelity (HF) PstI and XhoI (NEB) before insertion of the amplified *GPR1* using NEBuilder^®^ HiFI DNA assembly Cloning kit. After transformation of the StuI-linearized CIp10 plasmid into the *gpr1Δ/Δ* strain, we performed selection with nourseothricin. Furthermore, a *GPR1*-mTurquoise2 fusion was created. Therefore the CIp10 plasmid was digested with HF PstI and XhoI (NEB) before insertion of the amplified Gpr1 and the mTurquoise2, from a CIp10 – mTurqouise2 plasmid(44), using NEBuilder^®^ HiFI DNA assembly Cloning kit.

We introduced single point mutants using the Q5^®^ Site-Directed Mutagenesis kit (NEB). Primers containing the mutation amplified the entire CIp10–*GPR1*–mTurquoise2 plasmid and were confirmed through sequencing. The correct constructs with the mutated *GPR1* were transferred to a backbone CIp10 vector to limit mutations in the backbone of the plasmid that could have been introduced during the whole plasmid amplification. Primers utilized for introducing specific mutations and the transfer of the construct are listed in supplementary table 2.

### Selection of strains with a double integration of CIp10 using quantitative PCR

Gpr1 – mTurquoise2 strains containing two integrations of the CIp10 plasmid showed higher brightness and a distinct localization pattern. Therefore, all strains containing a CIp10 construct were selected for double integration using quantitative PCR. DNA was extracted using phenol/chloroform/isoamyl alcohol (PCI) followed by a double ethanol precipitation. DNA was diluted to 0.5 ng/μL and checked for a single genomic DNA integration of the plasmid using quantitative PCR. 2.5 ng/μL of DNA was utilized in the standard protocol of Promega with GoTaq^®^ qPCR Master Mix, CXR dye and primers listed in supplementary table 2 at a final concentration of 1 μM on the StepOnePlus Real-Time PCR system from Thermo Fisher Scientific. QPCR analysis was performed with Biogazelle Qbase+ software.

### Morphogenesis

A single colony was picked from each strain and grown overnight on synthetic complete medium with pH 5.5 containing 0.17% (w/v) yeast nitrogen base without amino acids, 0.5% (w/v) ammonium sulfate, 0.079% (w/v) complete supplement mix from MP biomedicals and 2% (w/v) glucose. Afterwards the cultures were diluted to O.D. 0,2 and grown for 4 hours till mid exponential phase. The O.D. of the resulting cultures were measured and diluted to achieve 100 cells/mL. 100 μL of the dilutions were distributed on plates containing synthetic minimal medium (synthetic complete medium at pH6 without complete supplement mix) with or without 100 μM methionine. Photos were taken after 4-5 days utilizing a Leica M165 C binocular microscope coupled with a Leica DFC420 C camera.

### β-glucan exposure quantification

The β-glucan exposure assay is based on the method reported by Ballou and colleagues (12). A single colony was grown overnight on synthetic complete medium containing 2% glucose before transferring to synthetic complete medium containing either 2% glucose or 2% glucose and 2% lactate at O.D. 0.2. After growing for 4 hours the number of cells in each culture was counted using a Guava^®^ flow cytometer and a volume containing 2.5 X 10^6^ cells was centrifuged and washed twice with FACS buffer containing 1% fetal bovine serum and 0,5mM EDTA in Phosphate buffered saline. After washing, the cells are resuspended in FACS buffer containing 5ng/μL Fc-hDectin1 (Invivogen) and incubated in the dark on ice for 1 hour. As a negative control we also included a sample which is not incubated with Fc-hDectin1. Subsequently, the cells were washed two times with FACS buffer to remove any nonbounded Fc-hDectin1 and resuspended in FACS buffer containing a 1:200 dilution of the secondary antibody F(ab’)2-Goat anti-Human IgG conjugated to AlexaFluor™ 488 (ThermoFisher Scientific). This solution was incubated for 1 hour on ice before finally washing twice with FACS buffer and measuring the fluorescence intensity of 5 000 cells using a Guava^®^ flow cytometer with 488nm excitation and fluorescence emission selected using a 522/30 bandpass filter.

### *C. albicans* survival within macrophages

We assayed the survival of *C. albicans* when challenged with bone marrow-derived macrophages (BMDMs) in an *in vitro* assay based on the report by Riedelberger and colleagues (45). BMDMs with a density of 1 x 10^5^ cells/well were infected with 1 x 10^4^ *C. albicans* cells in a 96-well plate and incubated for 3 hours at 37°C. To each well, 4% Triton-X 100 in PBS was added and the cells were dislodged from the well bottom. Cell suspensions were plated and colony forming units (CFU) were counted. Each assay was performed thrice with the same biological transformant due to practicalities.

### Assessing growth of *C. albicans* strains in macrophage differentiation medium

Growth of the *C. albicans* strains, which were utilized in the macrophage challenge assay, was assessed in the macrophage differentiation medium. Single colonies were grown overnight on synthetic complete medium before being washed twice with PBS and diluted to an O.D. of 0,05 in differentiation medium and distributed in a flat bottom 96-well plate. Growth was assessed by measuring the absorbance at 595nm every hour for 30 hours.

## Acknowledgments

This work was supported by the Research Foundation Flanders (FWO Vlaanderen) [1S01817N to W.V.G.] and the Research Council of the KU Leuven [3E210681 to W.V.G.]. We thank Martine De Jonge for the generous technical assistance in creation of the strains and performing the morphogenesis and β-glucan experiments.

