## Supplementary information for "Differential sensing by the *C. albicans* Gpr1 receptor results in morphogenesis, β-glucan masking and survival in macrophages"

**This PDF file includes:**

Supporting text
Figures S1 to S5
Tables S1 to S3

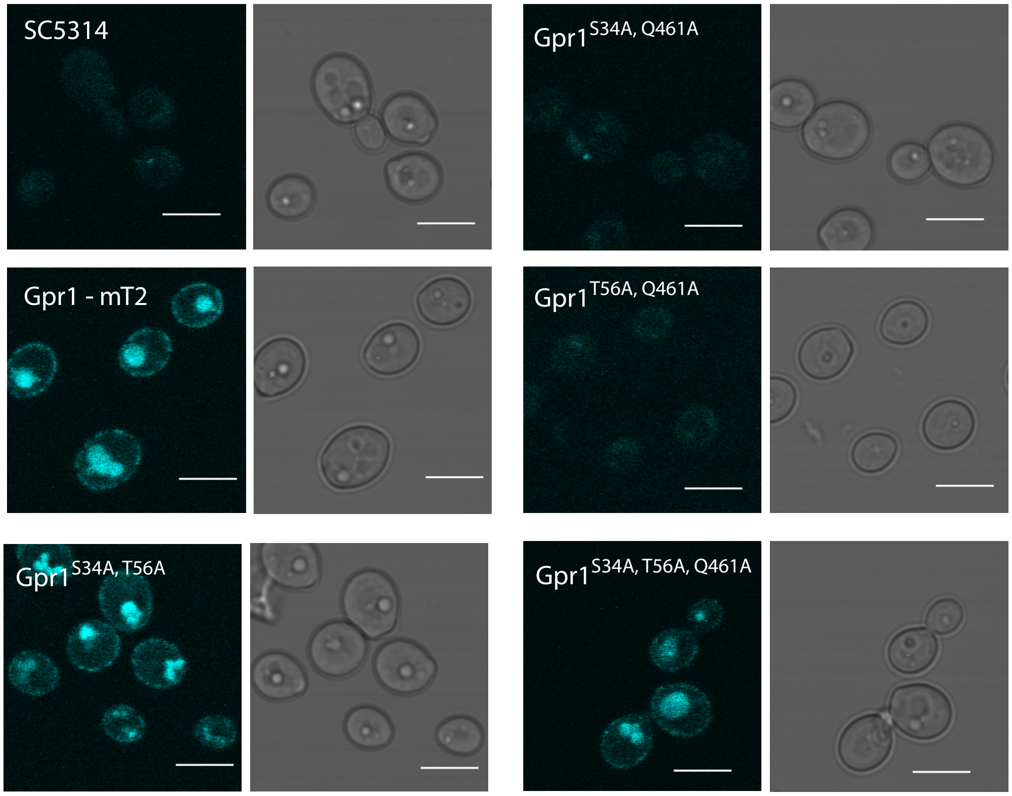


**Fig. S1**. Point mutations in Gpr1 at S34 and T56 does not hamper localization. Confocal fluorescence microscopy of mutated Gpr1 fused with mTurquoise2. mTurquoise2 was excited with 458nm light from an argon laser and emission was recorded through a bandpass filter BA480–495Scalebar is 5 μM.

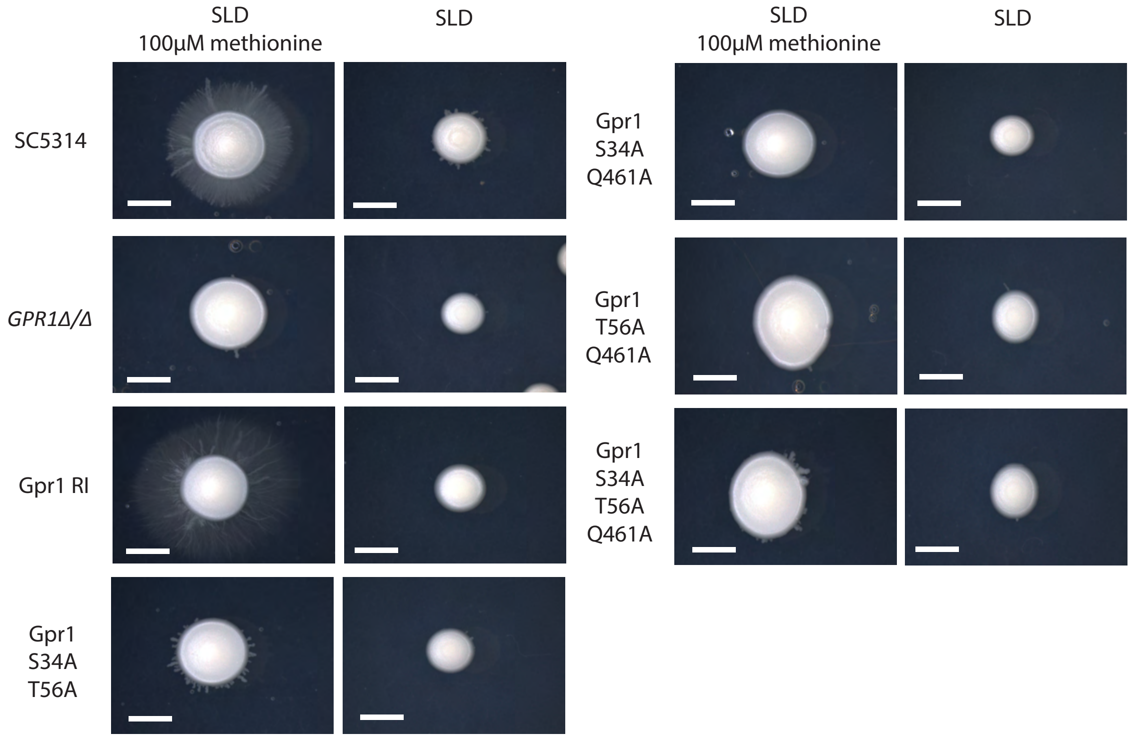


**Fig. S2.** Double and triple point mutants of Gpr1 confirm that the N-terminal tail is required for the methionine-induced morphogenesis. Morphogenesis assay on solid medium with and without 100μM L-methionine. Direct comparison of the SC5314 and GPR1 deletion strain with the double and triple Gpr1 point mutants. A representative colony is depicted for each strain in each condition. The experiment was performed twice with three biological repeats. Scalebar denotes 5 mm.

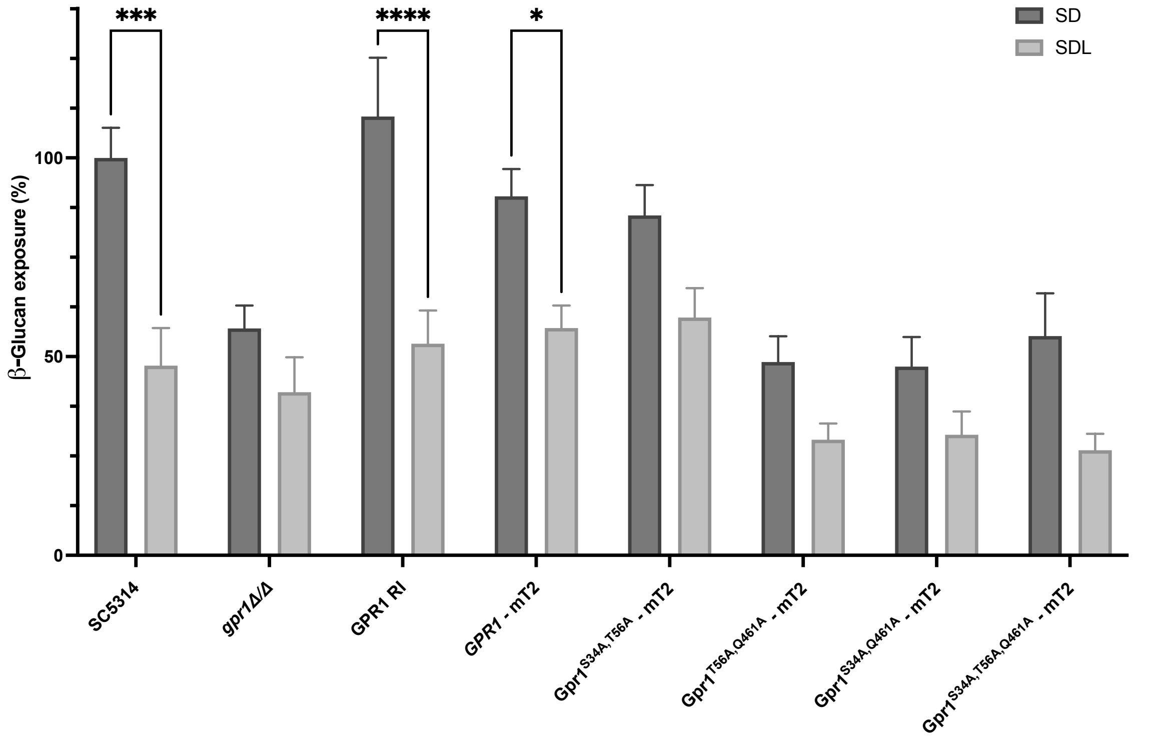


**Fig. S3** Double and triple point mutants of Gpr1 show intermediate and mixed glucan exposure phenotypes. Exposure of β-glucan as measured with flow cytometry after labeling with alexa488 utilizing hDectin1. Median fluorescence intensity of 5 000 cells is normalized to SC5314 in SD conditions. Statistical analysis was performed using two-way ANOVA with Sidak multiple comparison correction. 3 biological repeats were utilized for each strain in each condition and the experiment was performed twice. Asterisks denote significance levels with; *, $P < 0.05$; **, $P < 0.01$; ***, $P < 0.001$; ****, $P < 0.0001$. Depicted error bars show the mean and SD.

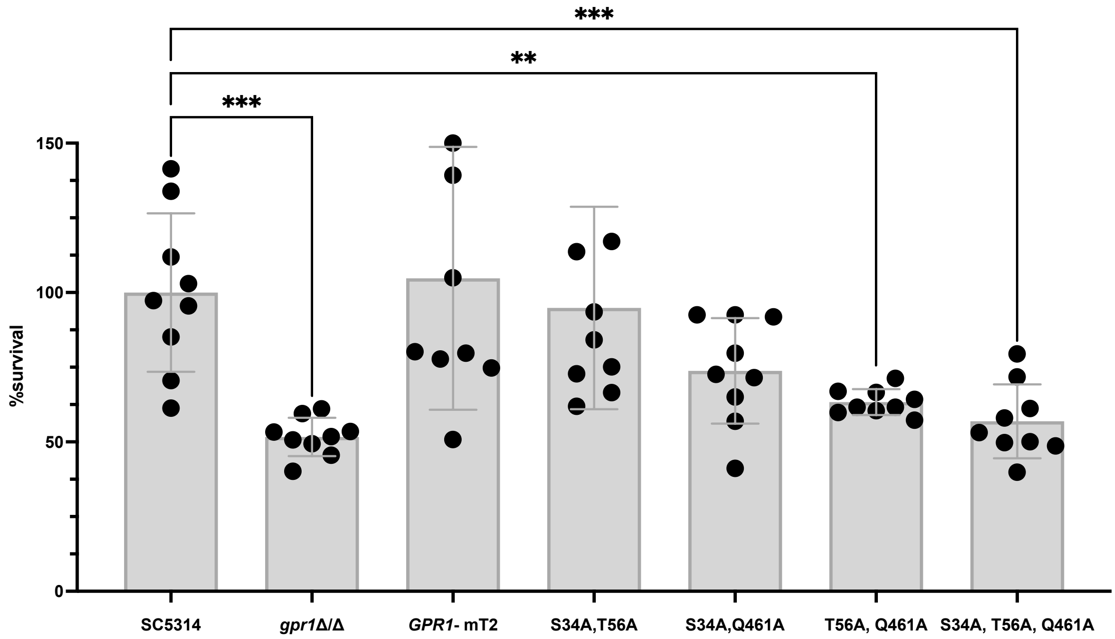


**Fig. S4** Double and triple point mutants of Gpr1 confirm the observation that Q461 has a role in survival when C. albicans is challenged with macrophages. Candida albicans survival assay when co-cultured with BMDMs for 3h at 37°C. Fungal cells were plated and counted to score for survival. Results shown are the average of three independent experiments containing three experimental repeats of a single biological repeat. Raw data is normalized to the survival rate of the wild type SC5314. Statistical analysis was performed using one-way ANOVA with a Bonferroni-corrected multiple comparison to the wild type. Asterisks denote significance levels with; *, $P < 0.05$; **, $P < 0.01$; ***, $P < 0.001$; ****, $P < 0.0001$. Depicted error bars show the mean and SD.

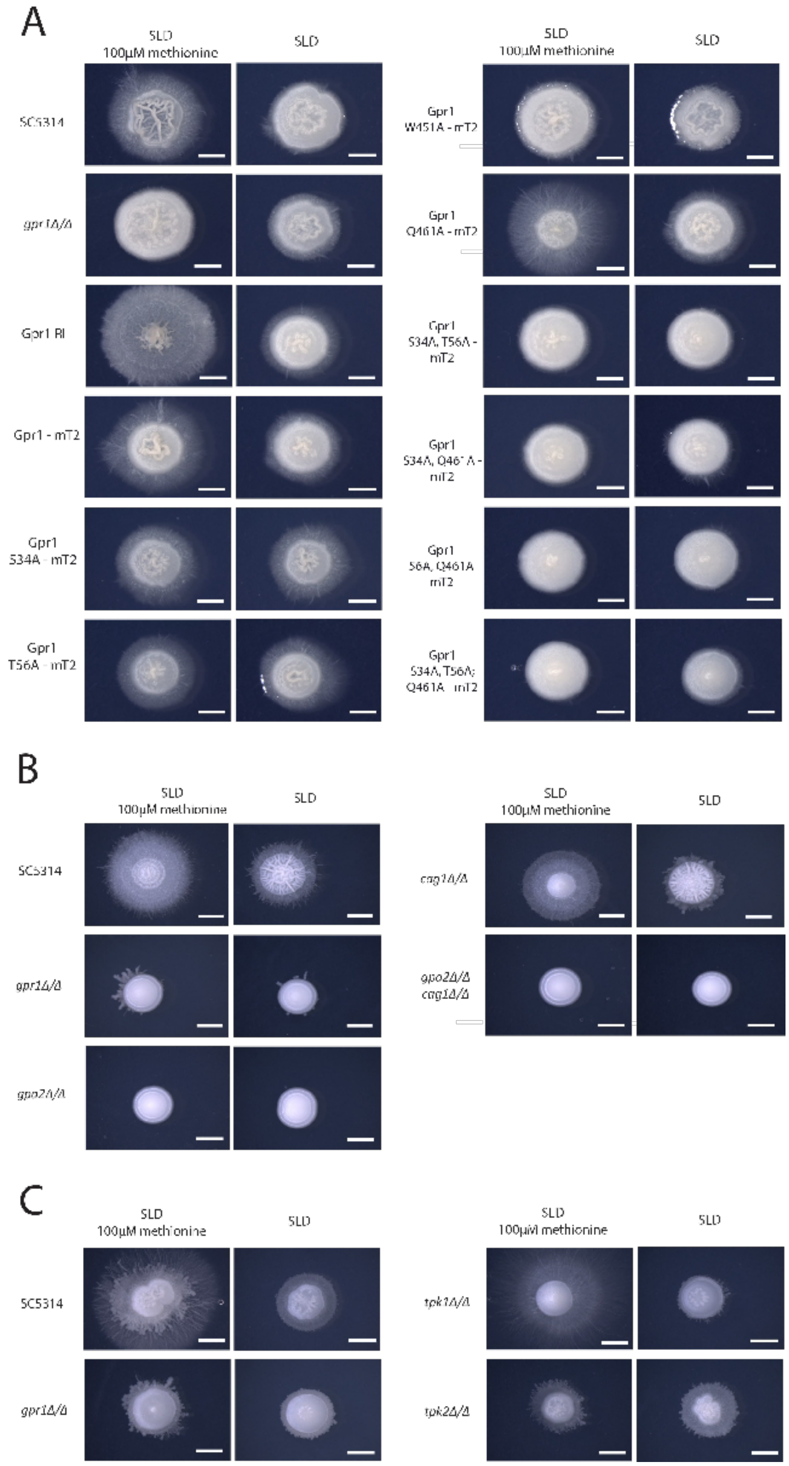


**Fig. S5**. The N-terminal tail is required for the methionine-induced morphogenesis through Gpa2 and Tpk2 at 37°C. Morphogenesis assay on solid medium at 37°C with and without 100µM L-methionine. Direct comparison of the SC5314 and GPR1 deletion strain with A) the double and triple Gpr1 point mutants, B) the putative G$\alpha$ subunits and C) the PKA catalytic subunits. A representative colony is depicted for each strain in each condition. The experiment was performed twice with three biological repeats. Scalebar denotes 5 mm. Panel A was color adjusted to fit panel B and C.

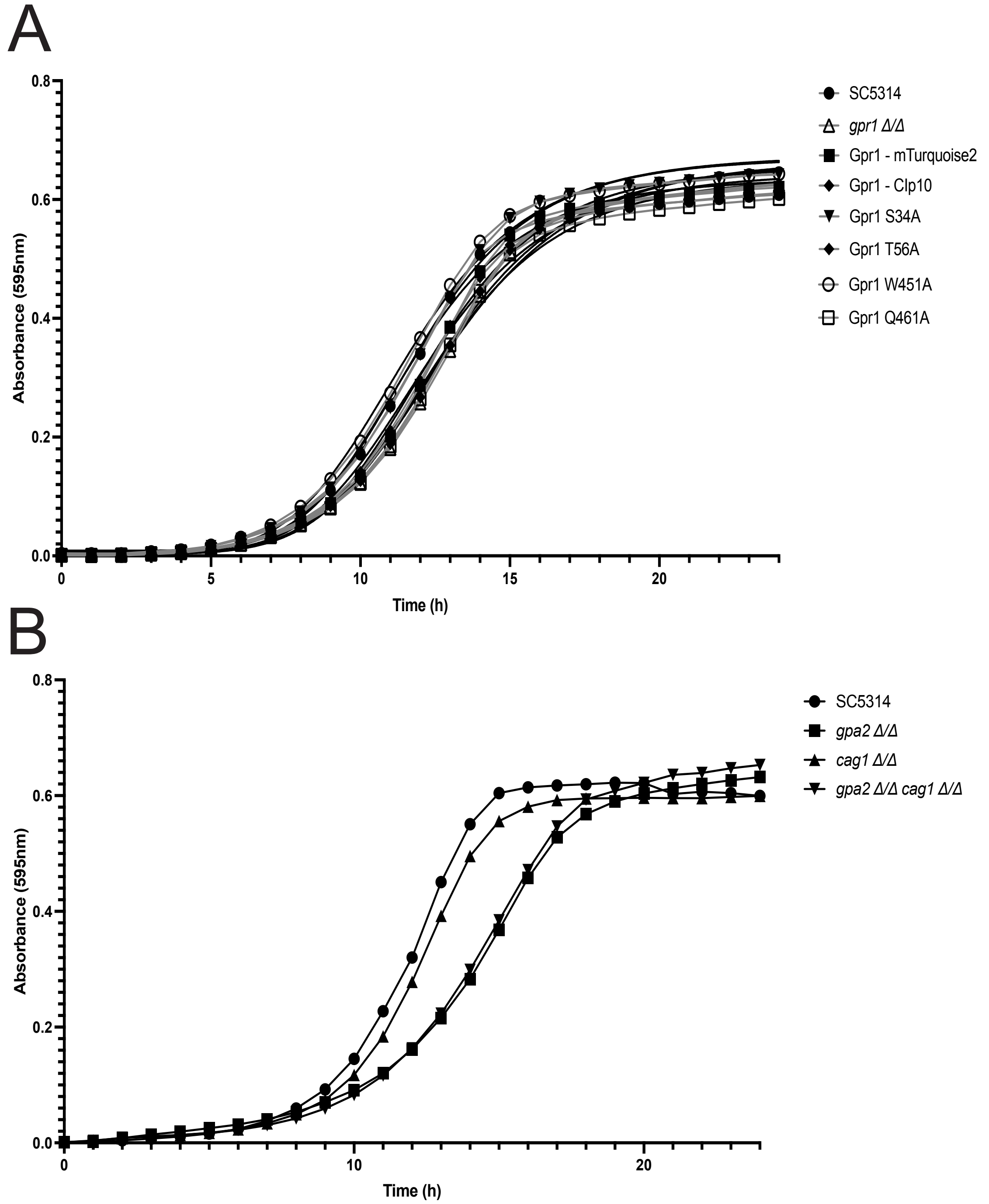


**Fig. S6** Growth curves of mutated Gpr1 strains and deletions of Gα proteins in macrophage differentiation medium. A) Deletion of GPR1 or mutation of Gpr1 at S34, T56, W451, Q461 into alanine does not result in significant growth defects. B) Disruption of the Gα proteins. Deletion of CAG1 does not result in a growth defect, whilst deletion of GPA2 results in a growth defect. Strains were diluted to an OD of 0,01 and grown in macrophage differentiation medium for 24 hours at 37°C. Depicted is the mean of each strain calculated from 3 biological repeats with 2 technical repeats. For clarity no standard deviation is depicted.

**Table S1.** List of strains utilized in this study

| Strain | Source | Identifier |
|---|---|---|
| SC5314 | ATCC | Cat#MYA-2876 |
| SC5314 – *gpr1Δ /gpr1Δ* | This study | WVG1 |
| SC5314 – *gpa2Δ /gpa2Δ* | This study | WVG2 |
| SC5314 – *cag1Δ /cag1Δ* | This study | WVG3 |
| SC5314 – *gpa2Δ /gpa2Δ - cag1Δ /cag1Δ* | This study | WVG4 |
| SC5314 -- *tpk1Δ /tpk1Δ* | This study | WVG5 |
| SC5314 – *tpk2Δ /tpk2Δ* | This study | WVG6 |
| SC5314 – *gpr1Δ /gpr1Δ* | This study | WVG7 |
| SC5314 – *gpr1Δ /gpr1Δ - RPS10-pACT1-GPR1-mTurquoise2-tACT1* | This study | WVG8 |
| SC5314 – *gpr1Δ /gpr1Δ - RPS10-pACT1-GPR1-mTurquoise2-tACT1* | This study | WVG9 |
| SC5314 – *gpr1Δ /gpr1Δ - RPS10-pACT1-GPR1-mTurquoise2-tACT1* | This study | WVG10 |
| SC5314 – *gpr1Δ /gpr1Δ - RPS10-pACT1-GPR1-mTurquoise2-tACT1* | This study | WVG11 |
| SC5314 – *gpr1Δ /gpr1Δ - RPS10-pACT1-GPR1-mTurquoise2-tACT1* | This study | WVG12 |

**Table S2.** List of primers utilized in this study

| Name | **Sequence (5'->3')** |
|---|---|
| **FW_Gpr1_ClP10** | TAATCATTCAAAATGCTGCACATGCCGGACCT AATATCAATAG |
| **FW_Gpr1_ClP10** | GATTTCCAGAATTTCACTCCCATTGGGGGTCC TTTTTTG |
| **RV_Gpr1_ClP10** | AATCATTCAAAATGCTGCACATGCCGGACCTA ATATCAATAG |
| **FW_Gpr1_mT2** | TAGAAACCATGCTAGCTGAACCACCTCC |
| **RV_Gpr1_mT2** | TTCAGCTAGCATGGTTTCTAAAGGTGAAG |
| **FW_mT2_Gpr1_Clp10** | GATTTCCAGAATTTCACTCCTTATTTGTACAAT TCATCCATAC |
| **RV_mT2_Gpr1_Clp10** | This study |
| **FW_S34A** | AATAGCTGCAgctGTATCTGCAG |
| **RV_S34A** | GTTGCTATAGTTGTCGGTATG |
| **FW_T56A** | TTCTACTACTgctGCATCTATACTTTCTG |
| **RV_T56A** | TCATCGGTGCTGTCCTTA |
| **FW_W451A** | TTGTCTTGTAgctCTTTTCCCATTTATTTTAC |
| **RV_W451A** | TAAGCAAATGGATAAATGAAAATC |
| **FW_Q461A** | ACAAGCCACCgctTTCAATTATGAAGAAG |
| **RV_Q461A** | AAAATAAATGGGAAAAGCC |
| **FW_Transfermutation_Clp10** | tagctgcatctgtatctgcaGCCACAGCAACAGTGACAA C |
| **RV_Transfermutation_Clp10** | acagtatgtaattcatcgatATCTTCATTGGATTTTACTC TATCAATTGG |
| **FW caAC1 promotor** | GGCTATGCCAATCAAAAAGG |
| **RV caACT1 promotor** | CCCCTTGGCCATAGGATATT |
| **TUB1_qPCR_FW** | TTACCCAGCTCCACAAGTGTC |
| **TUB1_qPCR_RV** | AAGTACAATCGGCGTGTTCC |
| **ADH1_qPCR_FW** | CACTCACGATGGTTCATTCG |
| **ADH1_qPCR_RV** | TAAGATTGGTGCGACATTGG |
| **18S_qPCR_FW** | GATGCCCTTAGACGTTCTGG |
| **18S_qPCR_RV** | CACGACGGAGTTTCACAAGA |

**Table S2.** List of plasmids utilized in this study

| Name | Source | Identifier |
|---|---|---|
| **pADH99** | Nguyen *et al.,* 2017 | pADH99 |
| **pADH110** | Nguyen *et al.,* 2017 | pADH110 |
| **Clp10-*GPR1*** | This study | pWVG1 |
| **Clp10-*GPR1*-mTurquoise2** | This study | pWVG2 |
| **Clp10-*GPR1*$^{S34A}$-mTurquoise2** | This study | pWVG3 |
| **Clp10-*GPR1*$^{T56A}$-mTurquoise2** | This study | pWVG4 |
| **Clp10-*GPR1*$^{W451A}$-mTurquoise2** | This study | pWVG5 |
| **Clp10-*GPR1*$^{Q461A}$-mTurquoise2** | This study | pWVG6 |
| **Clp10-*GPR1*$^{S34A,T56A}$-mTurquoise2** | This study | pWVG7 |
| **Clp10-*GPR1*$^{S34A, Q461A}$-mTurquoise2** | This study | pWVG8 |
| **Clp10-*GPR1*$^{T56A, Q461A}$-mTurquoise2** | This study | pWVG9 |
| **Clp10-*GPR1*$^{S34A, T56A, Q461A}$-mTurquoise2** | This study | pWVG10 |
